# Dual-layer transposon repression in heads of Drosophila *melanogaster*

**DOI:** 10.1101/267039

**Authors:** Marius van den Beek, Bruno da Silva, Juliette Pouch, Mohammed el amine Ali Chaouche, Clément Carré, Christophe Antoniewski

## Abstract

piRNA-mediated repression of transposable elements (TE) in the germline limits the accumulation of heritable mutations caused by their transposition in the genome. It is not clear whether the piRNA pathway plays a functional role in adult, non-gonadal tissues in *Drosophila melanogaster*. To address this question, we first analyzed the small RNA content of adult *Drosophila melanogaster* heads. We found that varying amount of piRNA-sized, ping-pong positive molecules in heads correlates with contamination by gonadal tissue during RNA extraction, suggesting that most of piRNAs detected in head sequencing libraries originate from gonads. We next sequenced the heads of wild type and *piwi* mutants to address whether *piwi* loss of function would affect the low amount of piRNA-sized, ping-pong negative molecules that are still detected in heads hand-checked to avoid gonadal contamination. We find that loss of *piwi* does not affect significantly these 24-28 RNA molecules. Instead, we observe increased siRNA levels against the majority of *Drosophila* transposable element families. To determine the effect of this siRNA level change on transposon expression, we sequenced the transcriptome of wild type, piwi, *dicer-2* and *piwi, dicer-2* double-mutant fly heads. We find that RNA expression levels of the majority of TE families in *piwi* or *dicer-2* mutants remain unchanged and that TE transcript abundance increases significantly only in *piwi, dicer-2* double-mutants. These results lead us to suggest a dual-layer model for TE repression in adult somatic tissues. Piwi-mediated transcriptional gene silencing (TGS) established during embryogenesis constitutes the first layer of TE repression whereas Dicer-2-dependent siRNA-mediated post-transcriptional gene silencing (PTGS) provide a backup mechanism to repress TEs that escape silencing by *piwi*-mediated TGS.

## Introduction

Transposable element (TEs) activity is thought to be an important force in genome evolution, as TE integration and excision can result in mutations that impact gene regulation networks (Tubio et al. 2014). However, these mutations may be detrimental to individuals, potentially decreasing lifespan and fertility (Wood et al. 2016). Therefore, limited TE mobilization is beneficial to both host and TE, whereas high TE activity decreases host fitness and adversely affects vertical transfer of the TE.

In *Drosophila melanogaster*, the siRNA (small-interfering RNA) and the piRNA (piwi-interacting RNA) pathways are important negative regulators of TE expression in somatic (Li et al. 2013; Ghildiyal et al. 2008a) and gonadal tissues (Vagin et al. 2006), respectively. Both pathways are active in the gonads, while the siRNA pathway is thought to be active in all somatic tissues. Dicer-2, the central endonuclease of the siRNA pathway operates on double-stranded RNA (dsRNA) molecules by processively translocating along the molecule and cutting every 21st nucleotide (Cenik et al. 2011; Welker et al. 2011). After the initial processing, these 21nt duplexes with 3’OH overhangs of 2 nt are rebound by Dicer-2 and r2d2 to be loaded into Argonaute-2 (Ago2). Ago2 can then engage in multiple rounds of endonucleolytic cleavage of transcripts with mature siRNA complementarity. Loss of functional siRNA pathway results in increased levels of TE expression and mobilization (Li et al. 2013; Xie et al. 2013; Czech et al. 2008; Ghildiyal et al. 2008a) and compromised male fertility as well as defective sperm development (Wen et al. 2015).

The *Drosophila* piRNA pathway is well characterized for its role in maintaining germline stem cells (Cox et al. 1998; Jin et al. 2013), genome integrity in the gonads and early embryo (Wang and Elgin 2011) and is therefore required for fertility. Its key components are 3 germline Argonaute-family proteins Piwi, Aubergine (Aub) and Argonaute-3 (Ago3). piRNA production relies on the processing of piRNA cluster transcripts, which are composed of TE fragments. The processing of these primary transcripts by the piRNA biogenesis machinery produces primary antisense piRNAs which are loaded into Piwi or Aub in a germline cells cytoplasmic structure called the *nuage* (Brennecke et al. 2007). Primary piRNA-loaded in Aubergine slices complementary sense TE transcripts between the 10^th^ and 11^th^ position, which will become the 5’ end of a new TE sense piRNA. Sense piRNAs are then loaded onto Ago3, which in turn slices complementary antisense piRNA cluster transcripts between the 10^th^ and 11^th^ position. This cytoplasmic cyclic process referred to secondary piRNA amplification leaves a detectable “ping-pong” signature, in which 5’ and 3’ ends of piRNAs tend to overlap by 10 nucleotides (Brennecke et al. 2007; Gunawardane et al. 2007).

piRNA-loaded Piwi proteins can re-enter the nucleus, where piRNAs guide Piwi towards complementary nascent transcripts. Piwi then recruits factors (Maelstrom, SuVar3-9, dSETDB1, HP1 and silencio/Panoramix) that establish and maintain H3K9me3 at the surrounding genomic vicinity of TE insertion sites, possibly including protein coding genes and hence functions in TGS (Sienski et al. 2012; Brower-Toland et al. 2007; Wang and Elgin 2011; Rangan et al. 2011; Sienski et al. 2015).

Zygotic piwi expression has been detected ubiquitously in early embryos up to the 14^th^ nuclear division (~2h after egg laying) (Mani et al. 2014; Rouget et al. 2010), depletion of Piwi in nurse cells and oocytes results in early arrest of embryonic development (Mani et al. 2014; Wang and Elgin 2011), and *piwi* acts as a suppressor of variegation in the eye (Gu and Elgin 2013; Pal-Bhadra et al. 2004), suggesting an important function for Piwi in germline cells maintenance and during early development of somatic tissues. In contrast the function of Piwi in larval or adult somatic tissue remains unclear. Piwi has both been reported to be present (Brower-Toland et al. 2007) or absent (Le Thomas et al. 2013) in 3^rd^ instar larval salivary glands whereas Aubergine and Ago3 have been observed in non-overlapping cells of the adult central nervous system (Perrat et al. 2013).

In order to unravel the role of piRNA pathway in TE control of somatic adult tissues, we analyzed small RNA profile in wild type, *piwi* and *dicer-2* mutant heads fly. We provide evidence that previously reported ping-pong pairs in adult heads likely result from contamination with testicular RNA, suggesting that secondary piRNA amplification does not take place in adult heads. However, small RNA sequencing of *piwi* mutant heads reveal an increased levels of siRNAs against most TE families. RNA-sequencing of *piwi* single mutant heads and *dicer-2* single mutant heads showed only minor upregulation of transposable elements, whereas double-mutants of *piwi* and *dicer-2* showed increased TE levels. Our results suggest a dual-layer model of TE repression in somatic tissues. The first layer of TE repression is established by Piwi at the chromatin level during early development as shown by Gu and Elgin (Gu and Elgin 2013). When TEs escape the epigenetic Piwi silencing, PTGS triggered by dicer-2 and siRNAs mediates TE degradation to decrease TE burden.

## Material and Methods

### Fly stocks

Flies were grown on standard *Drosophila* food at 25°C. All flies were brought into the w^m4^ background (Muller 1930). dicer-2^R416X^ and dicer-2^L811fsx^ alleles were previously described (Lee et al. 2004). piwi^2^ and piwi^3^ alleles were previously described (Cox et al. 1998). Double-mutants were generated by crossing virgin female dcr2^R416X^/CyO-GFP to male piwi^3^/CyO-GFP flies. Offspring virgin dcr2^R416X^/piwi^3^ flies were then crossed to male wm4;Ln^2R^ Gla, wgGla^1^, Bc^1^/ CyO-GFP to establish wm4;piwi^2^, dicer-2^R416X^/CyO-GFP stocks. Stocks were then screened by PCR for the presence of the *piwi* mutation. The same procedure was applied to generate wm4;piwi^3^, dicer-2^L811Fsx^/CyO-GFP stocks. The table below provides the detailed genotype of all mutant combinations used.

| Indicated name | Genotype | Maternal genotype | Paternal genotype |
| --- | --- | --- | --- |
| +/+ | *wm4*;+/+ | *wm4*;+/+ | *wm4*;+/+ |
| *dicer-2* G31R/- | *wm4*;dcr2<sup>G31R</sup>/dcr2<sup>L811Fsx</sup> | *wm4*;dcr2<sup>L811Fsx</sup>/CyO-GFP | *wm4*;dcr2<sup>G31R</sup>/CyO-GFP |
| *dicer-2* +/- | *wm4*;dcr2<sup>L811Fsx</sup>/+ | *wm4*;dcr2<sup>L811Fsx</sup>/CyO-GFP | *wm4*;+/+ |
| *dicer-2* -/- | *wm4*;dcr2<sup>L811Fsx</sup>/dcr2<sup>R416X</sup> | *wm4*;dcr2<sup>L811Fsx</sup>/CyO-GFP | *wm4*;dcr2<sup>R416X</sup>/CyO-GFP |
| *piwi* +/- | *wm4*;piwi<sup>2</sup>/+ | *wm4*;piwi<sup>2</sup>/CyO-GFP | *wm4*;+/+ |
| *piwi* -/- | *wm4*;piwi<sup>2</sup>/piwi<sup>3</sup> | *wm4*;piwi<sup>2</sup>/CyO-GFP | *wm4*;piwi<sup>3</sup>/CyO-GFP |
| *dicer-2* -/-, *piwi* -/- | *wm4*;piwi<sup>2</sup>, *dicer-2*<sup>R416X</sup>/piwi<sup>3</sup>,<br>dicer-2<sup>L811Fsx</sup> | *wm4*;piwi<sup>2</sup>,<br>dicer-2<sup>R416X</sup>/CyO-GFP | *wm4*;piwi<sup>3</sup>,<br>dicer-2<sup>L811Fsx</sup>/CyO-GFP |

### RNA extraction and sequencing

Head RNAs were prepared as followed. One-to two-day old flies were CO2 anesthetized, sorted by sex and genotype, transferred into 15 ml Falcon tubes and frozen in liquid nitrogen. The procedure was repeated multiple days until pools of 50 to 100 flies were obtained per biological replicate. Heads were separated from bodies by vortexing, followed by sieving and a careful selection of heads from thoraces and legs on a cooled metal plate. Heads were collected into 2 ml Precellys tubes for hard tissues and covered by 1ml Trizol. Heads were homogenized in two rounds of 5000 rpm for 30 seconds using a Precellys24 Tissue Homogenizer. Homogenate was centrifuged for 30 seconds at 13000 rpm and supernatant transferred into a new 2 ml tube, 200μl of Chloroform was added and tubes were thoroughly vortexed. Further purification was as in (Rio et al. 2010). Remaining DNA was removed using Fermentas DNase I, RNase-free following the manufacturers instructions.

Small RNA library preparation and sequencing was performed on an Illumina HiSeq 2500 at Fasteris Life Sciences SA (Plan-les-Ouates, Switzerland) using the *Drosophila* small RNA track based on the Illumina TruSeq protocol.

RNA-seq was performed in biological triplicates for +/+, *piwi* −/+, *piwi* −/−, *dicer-2* −/+ and *dicer-2* −/−, with one replicate per condition sequenced in paired-end mode (2 * 101) and two replicates sequenced in single-read mode (1*51). *piwi* −/−, *dicer-2* −/− samples were sequenced in biological duplicates in single-read mode. Total RNA was depleted of rRNA using Ribo-Zero™ Gold Kit (Epicentre). Directional RNA-seq library preparation and sequencing was performed at the Genomic Paris Centre (Paris, France) using the Epicentre ScriptSeq™ v2 RNA-Seq Library Preparation Kit on an Illlumina HiSeq 2000 instrument.

All sequence files generated in this study are available from the EBI European Nucleotide Archive database (http://www.ebi.ac.uk/ena/) under study accession numbers PRJEB8519 (small RNA-Seq GZI-2) and PRJEB25033 (other sequence runs).

### Computational Analysis

The complete computational analysis pipeline was run on our in-house Galaxy server. All necessary workflows and tools will be publicly available at http://mississippi.fr/

Histories and Figures can be reach using the following individual links.

Figures: 1A and 1B; 1C and 1E; 1D and 1F; 1G

Figures: 2A, 2B and 2C; 2D and 2E

Figures: 3A; 3B; 3C

Supplemental Figures: 1A and 1B; 1C; 1D; 1E; 1F

Supplemental Figures: 2A; 2B

**Figure 1.**
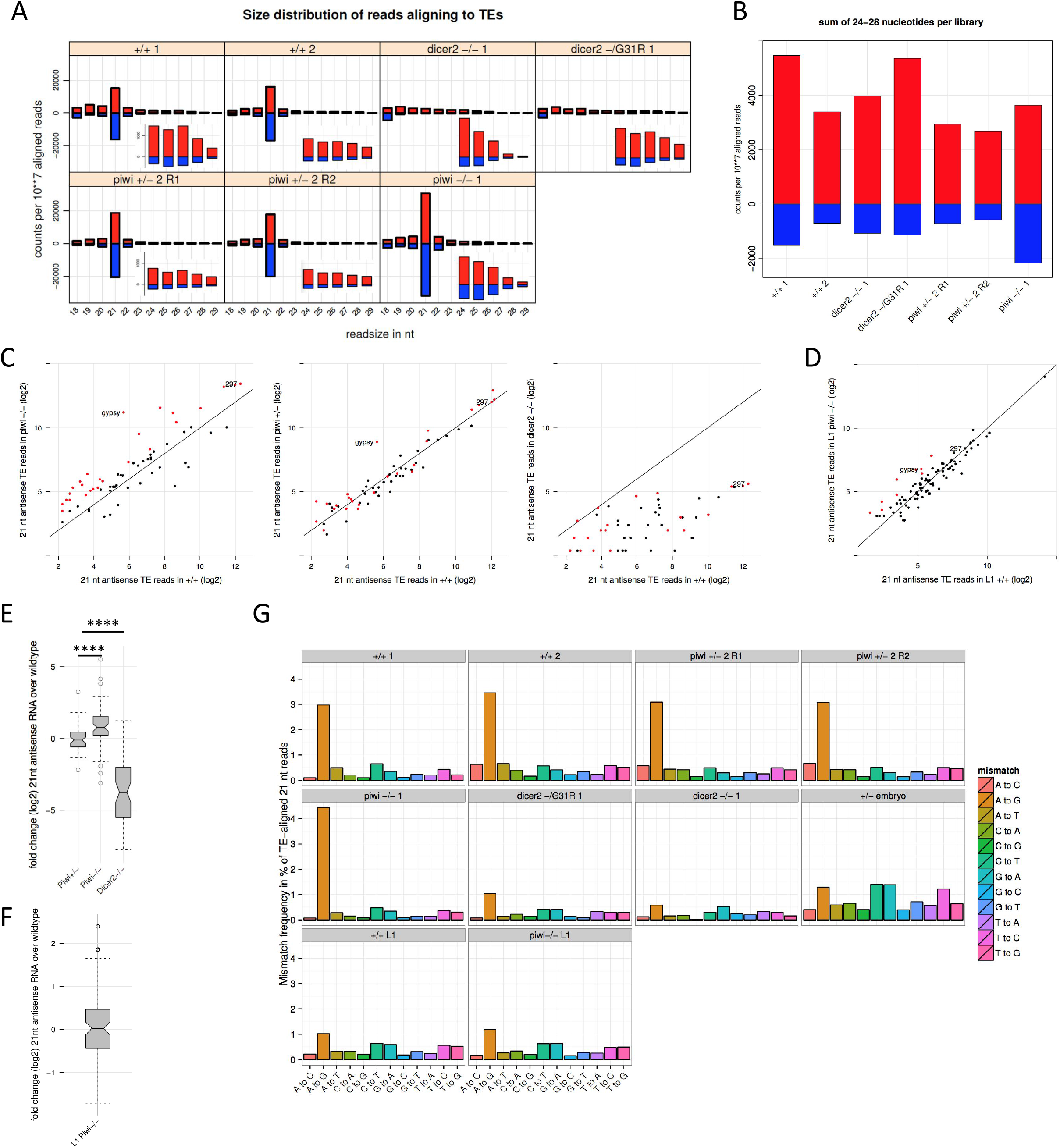
Loss of Piwi repression leads to an increase of TE-targeting siRNAs. (A) Overview of the size (x-axis) and amount (y-axis, in counts per 10 million mapped reads) of small RNA reads that align to TEs in adult heads of the indicated genotype. Zoom of the 24-28 nt fractions are shown as insets. (B) Sense or antisense reads of 24 to 28 nt were summed for each indicated genotype (x-axis). (C) Scatterplots displaying the abundance of 21 nt antisense reads in mutant (y-axis) and wild type (x-axis) heads. Red dots in the first panel indicate the transposon-specific 21 nt antisense reads that increased more than 2 fold in *piwi* homozygous mutant heads. These dots are shown for comparison in the second and third panel. (D) same as (C) but for wildtype 1^st^ instar larvae (homozygous *piwi* mutant *vs* wild type). (E) Boxplots showing the distribution of 21 nt antisense fold changes (y-axis) between wild type and the indicated mutants (x-axis). Significance of differences between the distributions was assessed with Mann-Whitney U test. (F) as (E), but for wildtype 1^st^ instar larvae (homozygous *piwi* mutant *vs* wild type). (G) Mismatches of 21 nt reads aligning to reference genome TE insertions with 1 mismatch allowed. Identity of mismatch is indicated on the x-axis, and the fraction of all reads with this mismatch identity over all TE matches is indicated on the y-axis. In all panels, the index of sample duplicate is indicated after the genotype when appropriate.

**Figure 2.**
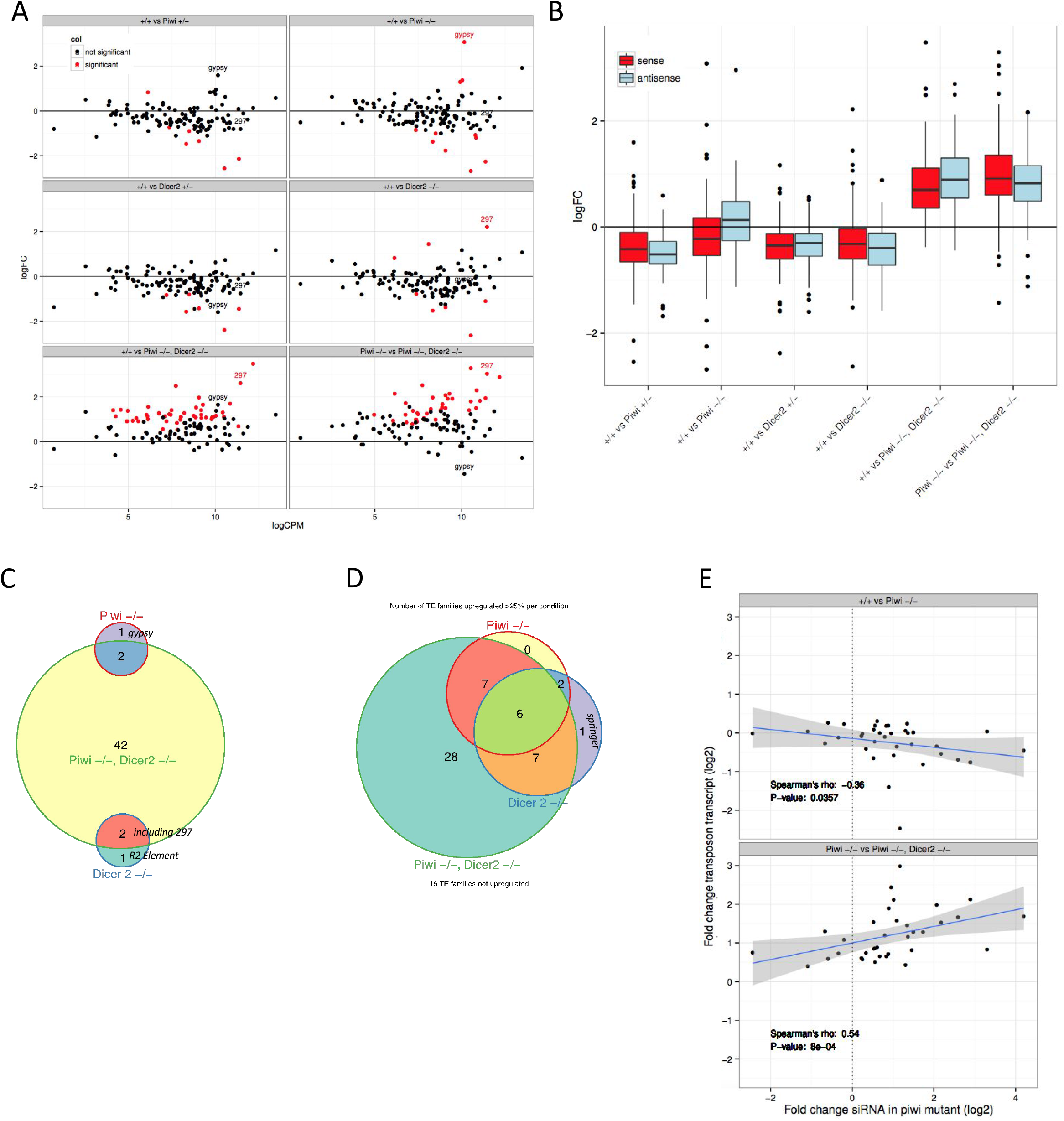
Piwi and Dicer-2 are complementary factors for the repression of TEs in adult heads. (A) Scatterplots displaying the log2 fold changes of sense TE transcript expression on the y-axis and the mean expression strength on the x-axis. log2 fold changes were calculated between the indicated mutant and the wild type TE levels, except in the last panel where we calculate changes between the double mutant and the *piwi* mutant. (B) Boxplots showing the distribution of log2 fold changes as in (A), but considering changes of sense (red) and antisense TE transcripts (blue) separately for each comparison (x-axis). (C) Venn Diagram showing the overlap of significantly (p.adj <0.05) up regulated TEs for the indicated mutants as compared to the wild type control. (D) as (C), but taking TEs whose transcript abundance increases more than 25% over the wild type. (E) Scatterplot displaying the correlation between log2 fold changes of 21 nt antisense RNA (siRNA) in *piwi* homozygous mutant heads compared to wild type heads on the x-axis and log2 fold changes of sense TE transcripts for the genotype comparisons indicated above each panel. Only TEs that passed a threshold of on average five 21 nucleotide antisense reads (after library-wise normalization, Material and Methods) over all small RNA libraries were analysed. Further, only TEs whose sense transcript level increased in *piwi, dicer-2* while insensitive to *piwi* loss are shown. A Scatterplot including all tested genotype that depicts all TEs that passed the siRNA threshold can be found in Supplemental Figure 2B. The blue line is a fit produced by the Imfit function and the grey area delimits the corresponding confidence interval.

**Figure 3.**
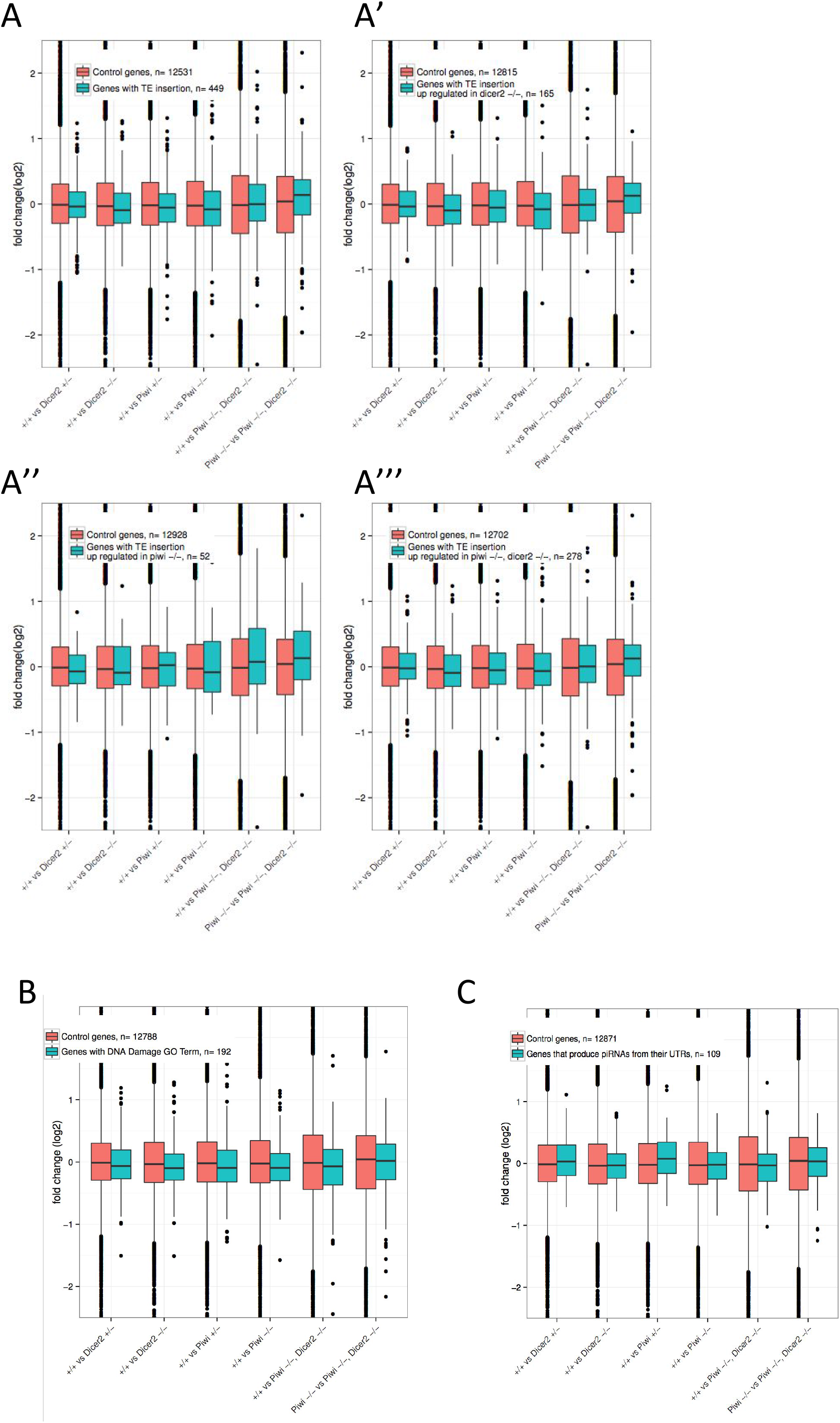
piwi, dicer-2 double mutant in heads is not affecting drastically previously reported piwi-regulated genes in gonads. Boxplots of fold change for genes with reference genome TE insertions in their genomic boundaries do not suggest a trend for these genes to become de-repressed across any of the tested mutant condition (x-axis). We compared genes without TE insertions (red boxes) with genes with insertion of TE of any family (blue boxes, A), or with TE insertions that are up regulated in *dicer2* homozygous (blue boxes, A’), *piwi* homozygous (A”) or *piwi, dicer-2* double homozygous mutants (A’“). Boxplots of the distribution of fold changes for genes with DNA Damage Gene ontology terms (B) and genes that have been reported to host piRNA production from their UTRs (C).

All small RNA libraries were quality controlled, sequencing adapter-clipped and converted to fasta reads. All reads that aligned to ribosomal RNA were discarded. All small RNA alignments were done using bowtie 0.12.7, allowing 1 mismatch between sequenced read and reference sequence (Langmead et al. 2009). To produce Supplemental Figure 1, fasta reads were aligned to the *Drosophila* genome (FlyBase release 5.49) (St Pierre et al. 2014), randomly placing reads that align equally well in multiple genomic locations (multimapper) using the bowtie option “-M 1”. Size distribution and ping-pong signature were calculated using the mississippi toolsuite (https://testtoolshed.g2.bx.psu.edu/view/drosofff/mississipi_toolsuite_beta). The ping-pong signature was calculated by counting the number of pairs that overlap between 5 to 15 nucleotides between sense- and antisense aligned reads and transforming the obtained counts into z-scores (each count subtracted by the mean and divided by the standard deviation). Ping-pong positive libraries were selected by having a z-score higher than 2 and more than 20 pairs overlapping by 10 nucleotides. Ping-pong negative libraries were selected by having a negative z-score. To obtain a list of differentially expressed miRNA between ping-pong positive and ping-pong negative libraries reads were matched to the *Drosophila* pre-miRNAs of the miRBase 20 release (Griffiths-Jones 2004; Griffiths-Jones et al. 2006, 2008; Kozomara and Griffiths-Jones 2011, 2014). Differential expression profiling between ping-pong positive and ping-pong negative libraries was performed using edgeR_3.8.2 (Robinson et al. 2010; McCarthy et al. 2012) with standard settings. For simulating contamination with testis RNA 2 testis-libraries (accessions SRX135547, SRX023726) were downsampled to 10 million reads, pooled and 50000 randomly selected reads were added to 2.45 *10^6^ randomly selected reads from ping-pong negative libraries. piRNA signature was calculated as before. Differential miRNA expression was calculated between simulated libraries and ping-pong negative libraries of equal size (randomly downsampled to 2.5 * 10^6^), with libraries that were sampled from the same initial ping-pong negative library paired as a blocking factor. This allows for obtaining an accurate list of miRNAs (contamination signature) that should be expected to be to significantly change in abundance if a contamination occurred. Size distribution for small RNAs that align to TEs (Fig. 1A, 1B) was calculated from reads that matched any of the canonical TE sequences with 1 mismatch allowed, excluding reads that matched to ribosomal RNA, tRNA or abundant insect viruses. Abundance of 21 nt antisense RNA for each TE family was calculated by filtering reads to 21 nt length and aligning reads to canonical TE sequences, allowing only unique reads using the bowtie option “-v 1”. Only antisense reads were counted, and only TEs with on average 20 reads per library were analyzed. Between-library normalized 21 nucleotide antisense TE counts were obtained by pooling these with miRNA reads (obtained as before) and calculating a normalization factor using the DESeq (Anders and Huber 2010) function estimateSizeFactors. log2 fold changes were calculated by dividing normalized reads of mutants by normalized reads of controls and taking the logarithm. Difference of the population of log2 fold changes were tested using a two-tailed Mann-Whitney-U test. To calculate mismatch frequencies for 21 nucleotide small RNA, ribosomal, non-coding RNA and viral reads were filtered out. Remaining reads were aligned to the collection of TE insertions (FlyBase version 5.49), allowing 1 mismatch. Each possible mismatch was counted and divided by the total number of 21 nucleotide reads aligned to the collection of TE insertions.

For gene expression profiling, reads were quality-filtered using the FASTX toolkit with a Quality cut-off of 30 for 90% of the read. For the paired-end libraries only the R1 read of the pair was used and trimmed to 51 nucleotides. Reads were then aligned to the *Drosophila* genome release 5 (dm3) using Tophat2 (Kim et al. 2013). Default parameters were used, except that we supplied Gene Model annotations from UCSC genome browser for dm3 (http://support.illumina.com/sequencing/sequencing_software/igenome.html). Read counting was performed using featureCounts (Liao et al. 2014) guided by the aforementioned Gene Model file.

For TE expression profiling, reads were further trimmed to 30 nucleotides and aligned to canonical TEs using bowtie 0.12.7, allowing 2 mismatches and only uniquely matching reads. Sense and antisense reads were counted and merge with gene counts. Differential expression profiling was performed using edgeR (Robinson et al. 2010; McCarthy et al. 2012). Genes with less than 5 reads on average across libraries were discarded from the analysis. Diverging from the default, we used Full Quantile between-library normalization as implemented by the EDAseq package (Risso et al. 2011) and removed unwanted variation using replicate samples with the RUVs function (choosing k=2) implemented in the RUVseq package (Risso et al. 2014). Library sequencing method (paired-end vs. single-read) was introduced together with gene-wise Full Quantile normalization offsets and gene-wise RUVs offsets as covariates in the edgeR design formula. All libraries were tested for differential gene expression against the wild type, and in addition the double-mutant was also tested against the piwi −/− mutant. Proportional Venn Diagrams in Figure 2C and Figure 2D were drawn using the Vennerable package (https://github.com/is229/Vennerable). The spearman rank correlation and corresponding P-value between log2 fold changes in TE 21 nucleotide antisense RNA (log2 fold change calculated from data underlying Figure 1C) and sense TE transcript expression was calculated with the rcorr function in the Hmisc R package (http://cran.r-proiect.org/web/packages/Hmisc/). All graphs were plotted using ggplot2 (http://ggplot2.org/). GO terms for DNA damage and genes with TE insertions were retrieved from FlyBase (St Pierre et al. 2014).

## Results

### Secondary piRNA biogenesis is not detectable in adult heads

Ago3 and Aubergine were detected by immunofluorescence in the optic lobe of the *Drosophila* central nervous system (Perrat et al. 2013) and several teams sequenced small RNA molecules in head samples which had the size (24-28 nt), 3’-end 2’-O-methylation and ping-pong signature of *Drosophila* piRNAs (Ghildiyal et al. 2008b; Yan et al. 2011a; Mirkovic-Hösle and Förstemann 2014a). However, these putative piRNAs corresponded to a very minor fraction of the sequence datasets, which raises the possibility of sample contamination by gonadal tissues during RNA extraction. To address this issue, we analyzed small RNA matching TE sequences in 24 small RNA sequencing libraries prepared from adult male heads isolated by sieving after freezing (Reinhardt et al. 2012). The majority of reads were in the size range of siRNAs (21 nt). However, we observed a fraction of reads in each library within the 24-28 nt, size range of *Drosophila* piRNAs (mean 7.5%, interquartile range 5.6-10.4%; Supplemental Fig 1A, 1B).

We searched for ping-pong partners among 24-28 nt reads for each of the 24 small RNA sequencing libraries, as had been done in previous works (Ghildiyal et al. 2008b; Yan et al. 2011a; Mirkovic-Hösle and Förstemann 2014a). We detected a significant ping-pong signature (>20 pairs, z-score of the 10 nt overlap >= 2) in 5 out of the 24 analyzed libraries (hereafter referred to as *ping-pong positive* libraries, Supplemental Fig 1C, red asterisks), while 7 libraries (hereafter referred to as *ping-pong negative* libraries, Supplemental Fig 1C, black square) had a negative 10 nt overlap z-score.

The genotype of any of the analyzed libraries was not expected to differentially affect piRNA biogenesis, suggesting that the significant signatures we found might be due to stochastic contamination by gonadal material during the RNA extraction from heads prepared from adult male isolated by sieving after freezing (Reinhardt et al. 2012). To investigate this possibility, we performed differential expression testing of miRNAs between the 5 ping-pong positive libraries and the 7 ping-pong negative libraries, under the assumption that RNA introduced from contaminating tissues would include tissue-specific miRNAs that are not normally expressed in heads. Under our assumption, those putative tissue-specific miRNAs would be absent in ping-pong negative heads libraries (not contaminated) and present in ping-pong positive heads libraries (contaminated).

A set of 27 miRNAs was significantly enriched in ping-pong positive libraries relative to ping-pong negative libraries at an adjusted P-value (Benjamini-Hochberg) of 0.01 (Supplemental Table S1). Since we analyzed male heads, the contaminating RNA giving rise to ping-pong signature might stem from the testis. To investigate this possibility we randomly added reads from testicular small RNA libraries (Toledano et al. 2012; Rozhkov et al. 2010) to ping-pong negative head libraries whose sizes were normalized by sampling to 2.5*10^6^ reads for comparison. We find that adding ~2% testes reads to mimic the contamination in the ping-pong negative head libraries is sufficient to detect a ping-pong signature (>20 pairs, z-score for 10 nt overlap >= 2; Supplemental Fig 1D).

In addition, we found a strong overlap of miRNAs differentially expressed between ping-pong positive and ping-pong negative libraries on the one hand, and miRNAs differentially expressed between ping-pong negative libraries with or without simulated testis contamination on the other hand: the 10 most significantly changed miRNAs between ping-pong negative libraries with or without 2% of testicular reads are also significantly changed between ping-pong positive and ping-ping negative libraries (9 miRNAs with p<0.01, 1 miRNA with p<0.025; Supplemental Table S1 and Supplemental Figure 1E). Altogether, these data strongly suggest that when detected in the head, the ping-pong signature is mostly due to contamination by gonadal abundant small RNAs during sample preparation, in particular if a sieving strategy after freezing is used to isolate the heads and not followed by eye check. However, we cannot exclude that Aubergine and Ago3 are producing secondary piRNAs in very low amounts or in a small set of head cells.

### Ping pong negative piRNA-like small RNAs in adult heads are not *piwi*-dependent

In agreement with our previous observations, new libraries prepared from wild type (+/+), heterozygous (*piwi* +/−) or homozygous (*piwi* −/−) piwi, and heterozygous (*dicer-2* +/−) or homozygous (*dicer-2* −/−) *dicer-2* heads carefully checked to avoid contamination by gonadal debris were ping-pong negative (Supplemental Figure 1F). However, we still detected a small amount of piRNA-sized reads (24-28nt) in these library (Figure 1A insets). Since Piwi is sufficient for piRNA biogenesis without ping-pong signature in ovarian follicle cells (Lau et al. 2009; Robine et al. 2009; Saito et al. 2009), we further explored the possibility that these 24-28nt piRNA-sized reads are Piwi-dependent primary piRNAs.

As the main function of Piwi has been proposed to induce transcriptional repression of TEs in ovaries (Sienski et al. 2012; Rozhkov et al. 2013), we focused on small RNA reads aligning to TEs.

We observed that for wild type, *piwi* heterozygous and *piwi* homozygous mutant heads, the 24-28 nucleotide reads show a slight bias for aligning to the sense strand of TEs (Figure 1A, inset), which is further amplified in *dicer-2* mutant flies. This is in contrast to piRNAs molecules in gonads that predominantly align to the antisense strand of TEs (Brennecke et al. 2007). Importantly, the fraction of 24-28 nucleotide antisense to TEs seemed to increase slightly in *piwi* homozygous mutant heads. Together, these results suggest that the small fraction of 24-28nt RNAs in ping-pong negative head libraries are not *piwi*-dependent RNAs (Figure 1B).

### Piwi mutation unmasks a TE-specific siRNA response in adult somatic tissues

We next examined whether loss of *piwi* would affect siRNAs that align TEs sequences in the head. In both wild type and heterozygous *piwi* mutant heads we detect a substantial amount of 21-nt sense and antisense reads, which increases in homozygous *piwi* mutant (Figure 1A, 1E). Importantly, these reads are strongly reduced in *dicer-2* mutants, confirming that TE-aligned reads are siRNAs (Figure 1A, 1E). We consistently observed the increase of 21 nucleotide sense and antisense RNAs in *piwi* homozygous mutant heads when considering only TE families that had on average more than 20 aligned reads per 10 million matched reads (“cp10m”). To quantify siRNA variations we only considered the fraction of 21 nucleotide reads that aligns to the TE complementary strand, in order to minimize quantification of partially degraded TE transcripts, which would be expected to align to the sense strand with little size specificity (discussed in (Malone et al. 2009)). We further restricted analysis to TE families that had on average five or more 21-nt antisense reads per library after applying a size factor to correct for sequencing depth. By doing so, we determined that in *piwi* mutants, 27 out of 62 transposable element families show >2 fold increases of siRNAs, with an overall median fold change of 1.69 compared to a 0.92 fold change for piwi heterozygous mutant heads (Figure 1C, P-value 1.03e-05, Mann-Whitney U). While most TE siRNAs were hence moderately increased, gypsy siRNA expression was increased more than 45 fold.

We also investigated if the observed changes would occur at earlier time points during development. We observed a 2.8 fold increase of gypsy-specific siRNA in *piwi^−/−^* mutant first instar larvae (L1) as compared to *piwi^+/+^* L1, however the majority of TE family siRNAs remained unchanged at this stage (p-value 0.83, Mann-Whitney U, Figure 1D).

Together, these results suggested that increased TE siRNA levels upon Piwi loss is a response that is strongest at the adult stage. In addition to the previously reported suppression of variegation of wm^4^ in adult eyes and mild decrease of HP1 occupancy at TEs loci in 3^rd^ instar larvae (Gu and Elgin 2013), loss of Piwi at early stage of development also induces an increased production of TE-specific siRNAs later on in adult somatic tissues.

### The siRNA response likely originates from Dicer-2-mediated TE transcript processing in adult heads

We next investigated the origin of the increase in TE derived siRNAs in *piwi* mutant heads. SiRNA response might be a direct consequence of increased TE transcription in the absence of Piwi, which in turn would result in increased processing of TE-derived dsRNAs by Dicer-2 into siRNAs. Alternatively, the elevated levels of TE-siRNAs might originate from piRNA clusters, which produce both piRNAs and siRNAs in the germline. If this were to be the case we should be able to detect an increase of specific 21-nucleotide RNAs originating from these piRNA clusters.

As most germinal piRNA clusters are transcribed bidirectionally we quantified both 21 nt sense and antisense RNAs that map exclusively to piRNA clusters (defined in (Brennecke et al. 2007)). We thereby excluded sequences shared with TE insertions elsewhere in the genome, allowing us to separate production of siRNAs originating from clusters and those originating from TE insertions. We observed a low quantity of 21nt reads derived from piRNA cluster in both *piwi* mutant and control wild type heads (between 2% and 3,5% of total TE reads, Supplemental Figure 2A). These cluster-derived 21 nt RNAs increased slightly in *piwi* mutant heads, although to a lesser extent that the increase of 21 nt antisense TE reads (1.44 fold vs 2.15 fold, Supplemental Figure 2B). The increase of TE-specific siRNAs in *piwi* mutant heads was thus unlikely to be caused by an increase in Dicer-2-mediated processing of piRNA cluster transcripts, favoring the hypothesis in which increased transcription of euchromatic TE insertions leads to an increase in TE-specific siRNA production through Dicer-2 activity. In principle, the increase of TE-specific siRNAs could be maternally inherited or stably maintained from early development. We took advantage of the fact that mature Ago2-loaded siRNAs are single-stranded and that double-stranded RNAs, among which the substrates of Dicer-2, are frequently deaminated through the action of Adenosine Deaminase Acting on RNA (ADAR) enzyme, which converts adenosine (A) to inosine (I) (Keegan et al. 2005; Palladino et al. 2000; Wu et al. 2011). This change manifests in a A to G mismatches in RNA sequencing datasets as compared to the DNA based reference genome. We therefore determined the frequency of all nucleotide mismatches for all FlyBase-listed TE insertions in our small RNA sequencing datasets (Figure 1G). The amount of A to G mismatches is not elevated over other mismatches in early embryos (1.5 % of all 21 nucleotide reads matching to TE insertions). We detect a similar frequency of A to G mismatches (1.2 to 1.4%) in first instar larvae, however no other mismatches were preferred, suggesting that ADAR might be active at low level in 1st instar larvae. In contrast, we detected a higher frequency of A to G mismatches in wild type heads (3.6 to 4.1%) that, despite a >2-fold increase of 21 nt RNA, further increases in *piwi* mutants (5.7%). Since this increase would not be observed if siRNA were inherited from earlier developmental stages, the data suggest that siRNA are actively produced by Dicer-2 from double-stranded TE RNA substrates in *piwi* mutant adult heads.

### Neither loss of Piwi nor Dicer-2 leads to strong upregulation of TEs

To determine whether the observed increase of siRNA production efficiently counteracts any increased TEs transcription caused by a lack of Piwi-mediated TGS at earlier stage, we sequenced the head transcriptome of *piwi* mutants, *dicer-2* mutants and *piwi, dicer-2* double-mutants and compared these to wild type head transcriptome. In *piwi* mutants, transcript levels of most TEs remain unchanged in those heads, with the notable exception of *gypsy*, whose level increases about 5-fold (Figure 2A). *Gypsy* is also the TE against which we observed the strongest increase of siRNA levels (Figure 1C), suggesting that the transcription of TEs is indeed increased in *piwi* mutant heads and correlates with Dicer-2 dependant siRNA production through the dicing of double-stranded TE RNAs.

Contrary to sense transposon transcripts, we did detect a significant increase in antisense transcripts in *piwi* mutant heads (Figure 2B, +/+ vs Piwi −/− blue boxes), that might form transient duplexes with sense TE transcripts, serving as a substrate for Dicer-2-mediated siRNA processing. Similarly, most TEs are not upregulated in *dicer-2* mutants, except for *297*, which produces a significant amount of siRNA in wild type heads, perhaps indicating inefficient Piwi-mediated TGS for this TE family.

We conclude that Piwi and Dicer-2 are redundant for the maintenance of TEs repression for most TE families in adult heads and that both Piwi-mediated TGS and Dicer-2-mediated PTGS can efficiently repress TEs.

### Piwi and Dicer-2 compensatory mechanism revealed in double-mutant

To confirm our hypothesis that the siRNA response in adult somatic tissues compensates for the loss of Piwi earlier in development, we analyzed RNA libraries from *piwi, dicer-2* double-mutant heads. We found that the majority of TE families is significantly upregulated in *piwi, dicer-2* double-mutant heads compared to single *piwi* or *dicer-2* mutants (Figure 2 A, B, C).

To illustrate the impact of the increased siRNA levels in *piwi* mutants on TE transcripts, we plotted for each TE family the log2 of the fold change of siRNAs in *piwi* mutants on the X-axis, and the fold change in TE transcript level on the y-axis. We see a tendency for TE transcripts that are targeted by more siRNAs in *piwi* mutants to decrease in abundance when comparing *piwi* mutant to the wildtype (Figure 2E, top panel). If we perform the same analysis but focus on the change of TE levels between *piwi* mutant and piwi, *dicer-2* double mutants (Figure 2E, bottom panel), we see a tendency for TEs transcripts to increase in abundance in the double mutant. This further suggests that increased siRNA production in *piwi* mutants efficiently counteracts loss of the transcriptional repression (TGS) established by *piwi* earlier during embryogenesis. If this compensation is failing in piwi, *dicer-2* double mutants TE transcript levels increase globally.

Altogether our results suggests a dual layer expression control of Piwi-mediated TGS and Dicer-2-mediated PTGS to firmly repress TEs levels in the adult soma.

### Piwi and Dicer2 do not restrict expression of genes containing TEs in their genomic loci

Piwi has previously been shown to silence genes adjacent to TEs insertions and genes that carry TEs sequences in their boundaries (Sienski et al. 2012). Our transcriptomic data does not support a significant trend of upregulation of genes that contain TE insertions in their genomic boundaries, whether we consider all TE families, or only those that are upregulated in piwi, *dicer-2* or piwi, *dicer-2* double-mutants (Figure 3A). Piwi has also been shown to repress genes that produce *traffic jam* class piRNAs from their 3’UTR (Robine et al. 2009). Again, we detect no expression bias for these genes in any of our heads mutant conditions (Figure 3C).

## Discussion

### Origin of piRNA-like molecules in somatic tissues

Through the analysis of a large number of small RNA sequencing libraries we revisited previous reports of putative secondary piRNA biogenesis in adult heads (Yan et al. 2011b; Ghildiyal et al. 2008a; Mirkovic-Hösle and Förstemann 2014b). Based on our analysis, it is likely that earlier observations resulted from small contamination with gonadal RNA, which can easily occur when collecting heads by vortexing frozen flies and filtering fly-parts by sieving. PiRNA-like molecules in adult *Drosophila* heads were previously observed in libraries that were beta-eliminated before sequencing, and/or that originated from siRNA pathway gene mutants (Ghildiyal et al. 2008; Mirkovic-Hösle and Förstemann 2014). Beta-elimination prevents sequencing of small RNAs with unprotected 3’OH groups thereby increasing the apparent sequencing frequency of 2’O-methylated contaminating piRNAs from gonadal origin. Likewise, decrease of siRNA sequencing in siRNA pathway mutant heads is expected to amplify the sequencing frequency of piRNAs, including contaminants from gonadal origin.

Our results highlight the importance of controlling the purity of RNA preparations. Considering the enormous increase of TE expression in piRNA mutant ovaries, often reaching more than 1000 fold upregulation, best practice for analysis should be a control that clearly shows the degree of tissue purity of the RNA preparation. We show that careful examination and hand-selection of fly heads eliminates most of artifactual detection of piRNA ping-pong patterns in head RNA preparations.

When contamination is limited, the marginal fraction of remaining 24-28 nt RNAs is not sensitive to Piwi loss. This implies that either these piRNA-like molecules are primary piRNAs produced independently of Piwi (Nagao et al. 2010) or that they are degradation products. Together with their sense bias, the absence of a marked peak in the piRNA-size range (Figure 1A) and the low transcriptional activity of piRNA clusters in heads, these observation strongly suggest that these 24-28 nt RNAs are not piRNAs.

### Piwi likely exerts its function in adult heads through inherited transcriptional repression set in early development

Zygotic p*iwi* mutations are suppressors of variegation, revealing a link between the piRNA pathway and heterochromatin silencing (Pal-Bhadra et al. 2004). In contrast, late inactivation of *piwi* in eye imaginal discs does not suppresses variegation (Gu and Elgin 2013), suggesting that Piwi silencing established during early embryogenesis is maintained in the absence of Piwi product. As we were unable to evidence a role of piRNAs in TE silencing in adult heads and found that zygotic loss of *piwi* alone has little effect on steady state RNA levels of most of TE families in heads, our work is in line with this model.

### Repression of TEs in the absence of siRNAs

*Dicer-2* mutations caused a strong reduction of TE-specific siRNAs. However, this did not result in major changes of TEs RNA expression for most TE families in adult heads, with the notable exception of *297*. This observation is in line with the results of Ghildiyal *et al* (Ghildiyal et al. 2008b) who found *297* expression to be strongly increased in heads of *dicer-2* mutants, and Xie *et al* (Xie et al. 2013) who demonstrated increased somatic transposition of 297 in *dicer-2* mutants. How repression of specific TEs escapes from Piwi and mainly relies on siRNA pathways requires further investigation.

### A dual-layer TEs repression by small RNAs

We show that in adult heads, the loss of only *piwi* or *Dicer-2* does not lead to significant change in expression of most of TE families. However, loss of piwi is associated to increased levels of antisense TE transcripts and of siRNAs, whereas double *piwi Dicer-2* mutants show increased levels of transcripts from a large panel of TE families. Similar observations were made recently in ovarian somatic cells where double knockdown of piwi and Dicer-2 leads to synergistic derepression of LTR retrotransposons (S. Chambeyron, personal communication). We thus propose a dual-layer repression mechanism whereby residual expression of TE that results from incomplete piwi-mediated TGS feeds Dicer-2 for siRNA production that in turn reduces TE transcript levels through PTGS. This mechanism might be especially relevant for TEs that can escape Piwi-mediated repression, such as 297, or TEs that do not induce yet Piwi-mediated TGS because they are not integrated into piRNA clusters. It could also play a role in aging since an increase in TE expression and transposition was observed in aging adult flies, which was amplified in *Dicer-2* mutants and mitigated by *Dicer-2* overexpression (Wood et al. 2016).

In evolutionary terms, the “failsafe”, dual-layer repression mechanism could help during the early steps of a TE invasion of a fly population. Thus, siRNA-mediated PTGS would maintains a tolerable load of the TE transcripts, until the TE has integrated into a piRNA cluster and triggered a stable Piwi-mediated TGS.

## Acknowledgements

We thank S. Chambeyron and Alain Pelisson for sharing unpublished data and the Galaxy community for their support. This work received financial support to CA from the Agence Nationale de la Recherche grants ANR-13-BSV2-0007 and ANR-10-BLAN-1210. MdvB had a PhD fellowship from the Ministère de la Recherche et de l’Enseignement Supérieur. The École normale supérieure genomic core facility was supported by the France Génomique national infrastructure, funded as part of the “Investissements d"Avenir" program managed by the Agence Nationale de la Recherche (contract ANR-10-INBS-09).

## Author Contributions

MvdB performed the experiments and the computational analyses and prepared figures and tables. JP and MeaAC performed the RNA sequencing experiments. BdS performed genetic crosses. CA conceived the project. MvdB, CC and CA wrote the manuscript.

---

**Supplemental Figure 1.**
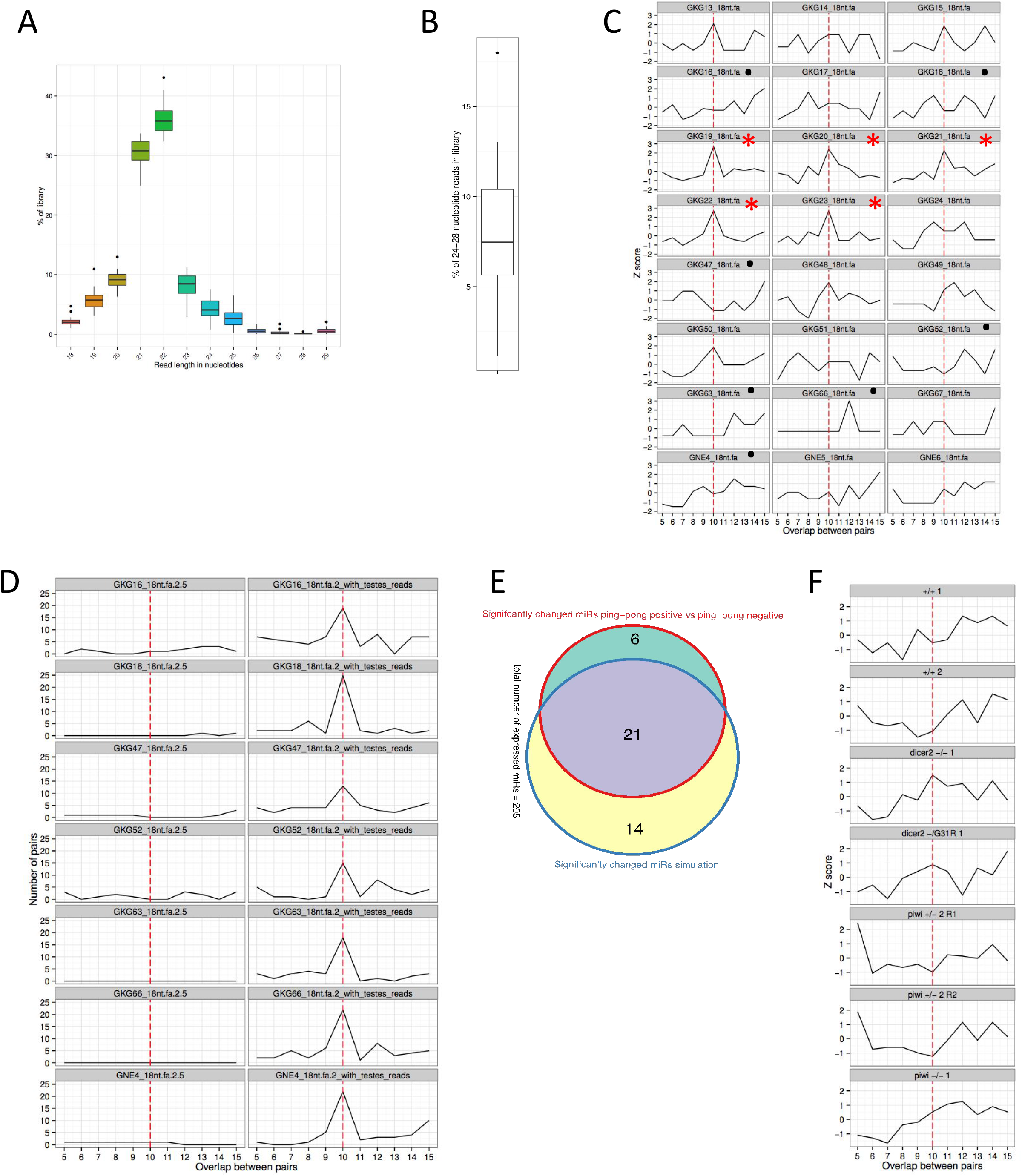
Drosophila small RNA head libraries have variable amount of piRNA-sized reads and ping-pong signature in heads, which correlates with a contamination signature. (A) Boxplot showing the distribution of read lengths for the libraries analyzed in (C). (B) similar to (A), but for the fraction of 24-28 nucleotide length reads. (C) Ping-pong signature - tendency for small RNAs to overlap. The number of overlaps was transformed to Z scores to take into account the “cleanness” of the 10 nucleotide overlap as compared to other lengths of overlaps. Libraries with red asterisks were selected as ping-pong positive, and black dots indicate libraries selected as ping-pong negative libraries. (D) Ping-pong signature for 2.5*10^6^ ping-pong negative libraries with (right group of panels) and without (left panels) the addition of 2% of a test¡s-l¡brary. Number of pairs are shown instead of Z scores, as some down-sampled libraries had 0 overlapping pairs. (E) Venn Diagram showing the overlap of differentially expressed miRNAs between ping-pong positive compared to ping-pong negative libraries (red circle) and ping-pong negative libraries compared to ping-pong negative libraries with the addition of 2% of a testis library (blue circle). (F) ping-pong signature for small RNA libraries (as in (C)) of indicated genotypes of heads closely dissected by hands.

**Supplemental Figure 2.**
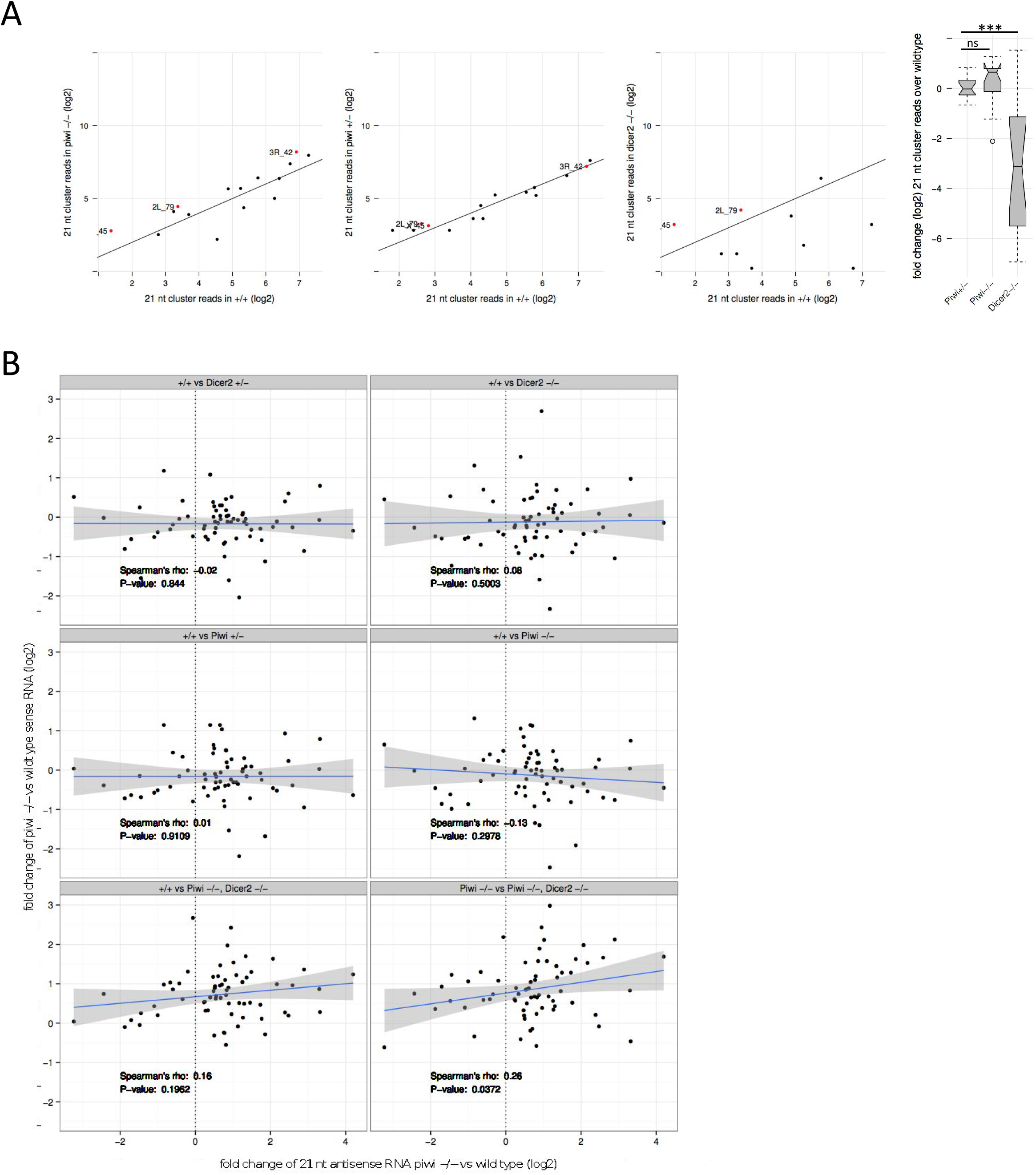
(A) Scatterplots displaying the abundance of cluster-derived 21 nt reads in mutant (y-axis) and wild type (x-axis) heads. Red dots in the first panel indicate the cluster-specific 21 nt antisense reads that increased more than 2 fold in *piwi* −/− mutant heads. These dots are shown for comparison in the second and third panel. (A-right) Boxplots showing the distribution of 21 nt read fold changes (y-axis) between wild type and the indicated mutants (x-axis). Significance of differences between the distributions was assessed with Mann-Whitney U test. (B) Related to Figure 2E. Scatterplot displaying the correlation between log2 fold changes of 21 nt antisense RNA in *piwi* homozygous mutant heads compared to wild type heads on the x-axis and log2 fold changes of sense TE transcripts for the genotype comparisons indicated above each panel. All TE families that passed a threshold of on average five 21 nucleotide antisense reads over all small RNA libraries were analysed. The blue line is a fit produced by the Imfit function, and the grey area delimits the corresponding confidence interval.

**Supplementary Table S1.** Differential expression testing using EdgeR between ping-pong negative and ping-pong positive libraries (Sheet #1) The top 10 differentially detected miRNA between ping-pong negative libraries and ping-pong negative libraries supplemented with 2% testicular reads (Sheet 2) are highlighted in red. Original data at https://lbcd41.snv.jussieu.fr/artbio/u/marius-ged/h/compare-simulation-with-real-differences

| Name | logFC | logCPM | LR | PValue | adj.p.value | Dispersion | Total reads |
| --- | --- | --- | --- | --- | --- | --- | --- |
| dme-mir-31b | 2.57E+00 | 6.10E+00 | 6.44E+01 | 1.03E-15 | 2.25E-13 | 1.34E-01 | 3.86E+03 |
| dme-mir-959 | 3.76E+00 | 3.61E+00 | 4.95E+01 | 1.98E-12 | 2.16E-10 | 3.19E-01 | 6.71E+02 |
| dme-mir-991 | 2.51E+00 | 3.22E+00 | 3.81E+01 | 6.59E-10 | 4.79E-08 | 1.85E-01 | 4.96E+02 |
| dme-mir-983-2 | 2.78E+00 | 2.65E+00 | 3.62E+01 | 1.75E-09 | 9.56E-08 | 2.18E-01 | 3.22E+02 |
| dme-mir-961 | 2.99E+00 | 2.69E+00 | 3.44E+01 | 4.40E-09 | 1.92E-07 | 2.85E-01 | 3.38E+02 |
| dme-mir-310 | 3.81E+00 | 5.06E-01 | 3.18E+01 | 1.68E-08 | 5.36E-07 | 1.80E-01 | 5.40E+01 |
| dme-mir-963 | 3.28E+00 | 2.01E+00 | 3.18E+01 | 1.72E-08 | 5.36E-07 | 3.20E-01 | 1.95E+02 |
| dme-mir-960 | 2.72E+00 | 4.18E+00 | 3.15E+01 | 2.04E-08 | 5.56E-07 | 2.97E-01 | 1.03E+03 |
| dme-mir-985 | 3.86E+00 | 3.66E+00 | 2.99E+01 | 4.65E-08 | 1.13E-06 | 5.82E-01 | 6.96E+02 |
| dme-mir-2494 | 4.26E+00 | 6.81E-02 | 2.79E+01 | 1.28E-07 | 2.65E-06 | 1.59E-01 | 3.20E+01 |
| dme-mir-983-1 | 2.48E+00 | 2.54E+00 | 2.78E+01 | 1.34E-07 | 2.65E-06 | 2.32E-01 | 2.92E+02 |
| dme-mir-982 | 1.88E+00 | 3.77E+00 | 2.63E+01 | 2.85E-07 | 5.00E-06 | 1.61E-01 | 7.42E+02 |
| dme-mir-312 | 2.18E+00 | 3.18E+00 | 2.63E+01 | 2.98E-07 | 5.00E-06 | 2.09E-01 | 4.84E+02 |
| dme-mir-iab-8 | 3.52E+00 | 5.98E-01 | 2.37E+01 | 1.12E-06 | 1.74E-05 | 3.05E-01 | 5.80E+01 |
| dme-mir-375 | 1.21E+00 | 9.09E+00 | 2.27E+01 | 1.94E-06 | 2.57E-05 | 9.16E-02 | 3.04E+04 |
| dme-mir-977 | 2.70E+00 | 3.42E+00 | 2.26E+01 | 1.97E-06 | 2.57E-05 | 4.01E-01 | 5.84E+02 |
| dme-mir-976 | 2.92E+00 | 5.33E-01 | 2.26E+01 | 2.00E-06 | 2.57E-05 | 1.86E-01 | 5.40E+01 |
| dme-mir-984 | 1.94E+00 | 2.56E+00 | 2.20E+01 | 2.73E-06 | 3.31E-05 | 1.86E-01 | 3.05E+02 |
| dme-mir-956 | 3.35E+00 | 8.13E+00 | 2.06E+01 | 5.75E-06 | 6.59E-05 | 7.09E-01 | 1.58E+04 |
| dme-mir-314 | 3.66E+00 | 7.04E+00 | 1.94E+01 | 1.04E-05 | 1.13E-04 | 8.74E-01 | 7.46E+03 |
| dme-mir-962 | 2.67E+00 | 5.17E-01 | 1.84E+01 | 1.79E-05 | 1.86E-04 | 2.14E-01 | 5.20E+01 |
| dme-mir-4966 | 2.82E+00 | 9.68E-02 | 1.74E+01 | 3.01E-05 | 2.92E-04 | 1.58E-01 | 3.30E+01 |
| dme-mir-974 | 1.84E+00 | 1.61E+00 | 1.74E+01 | 3.08E-05 | 2.92E-04 | 1.70E-01 | 1.42E+02 |
| dme-mir-1015 | 2.91E+00 | 3.39E-01 | 1.50E+01 | 1.06E-04 | 9.62E-04 | 3.74E-01 | 4.50E+01 |
| dme-mir-311 | 2.16E+00 | 3.77E+00 | 1.48E+01 | 1.19E-04 | 1.04E-03 | 4.11E-01 | 7.50E+02 |
| dme-mir-4914 | 2.90E+00 | -2.59E-01 | 1.32E+01 | 2.84E-04 | 2.38E-03 | 1.62E-01 | 2.10E+01 |
| dme-mir-973 | 2.03E+00 | 8.04E-01 | 1.09E+01 | 9.60E-04 | 7.75E-03 | 3.15E-01 | 6.90E+01 |
| dme-mir-iab-4 | 1.57E+00 | 1.27E+00 | 1.01E+01 | 1.52E-03 | 1.18E-02 | 2.07E-01 | 1.05E+02 |
| dme-mir-958 | 2.76E+00 | 3.83E+00 | 9.00E+00 | 2.69E-03 | 2.02E-02 | 1.12E+00 | 7.52E+02 |
| dme-mir-964 | 1.51E+00 | 1.19E+00 | 8.60E+00 | 3.37E-03 | 2.45E-02 | 2.37E-01 | 1.00E+02 |
| dme-mir-4983 | -3.25E+00 | -3.19E-01 | 8.34E+00 | 3.88E-03 | 2.73E-02 | 7.58E-01 | 1.90E+01 |
| dme-mir-4939 | 2.81E+00 | -5.69E-01 | 7.97E+00 | 4.75E-03 | 3.24E-02 | 3.28E-01 | 1.20E+01 |
| dme-mir-12 | 5.09E-01 | 1.12E+01 | 7.47E+00 | 6.29E-03 | 4.16E-02 | 4.93E-02 | 1.31E+05 |
| dme-mir-979 | 2.33E+00 | -4.59E-01 | 7.15E+00 | 7.50E-03 | 4.81E-02 | 2.37E-01 | 1.50E+01 |
| dme-mir-33 | -7.46E-01 | 1.19E+01 | 6.91E+00 | 8.58E-03 | 5.35E-02 | 1.09E-01 | 2.04E+05 |
| dme-mir-34 | -4.93E-01 | 1.54E+01 | 6.69E+00 | 9.71E-03 | 5.88E-02 | 4.97E-02 | 2.29E+06 |
| dme-mir-972 | 1.18E+00 | 2.38E+00 | 6.31E+00 | 1.20E-02 | 7.08E-02 | 2.61E-01 | 2.63E+02 |
| dme-mir-31a | 5.52E-01 | 1.19E+01 | 6.02E+00 | 1.42E-02 | 8.13E-02 | 7.21E-02 | 2.09E+05 |
| dme-mir-10 | 5.94E-01 | 1.12E+01 | 5.90E+00 | 1.52E-02 | 8.32E-02 | 8.49E-02 | 1.29E+05 |
| dme-mir-978 | 2.48E+00 | -6.88E-01 | 5.89E+00 | 1.53E-02 | 8.32E-02 | 2.90E-01 | 9.00E+00 |
| dme-mir-92a | -1.01E+00 | 6.43E+00 | 5.82E+00 | 1.59E-02 | 8.43E-02 | 2.31E-01 | 4.92E+03 |
| dme-mir-4964 | 1.74E+00 | -3.44E-01 | 5.78E+00 | 1.62E-02 | 8.43E-02 | 9.98E-02 | 1.80E+01 |
| dme-mir-4972 | 2.73E+00 | -9.28E-01 | 5.41E+00 | 2.00E-02 | 1.01E-01 | 3.22E-01 | 4.00E+00 |
| dme-mir-966 | -5.83E-01 | 5.34E+00 | 5.31E+00 | 2.13E-02 | 1.05E-01 | 8.08E-02 | 2.05E+03 |
| dme-mir-100 | -6.04E-01 | 9.61E+00 | 5.21E+00 | 2.25E-02 | 1.09E-01 | 9.50E-02 | 4.20E+04 |
| dme-mir-184 | 6.53E-01 | 1.52E+01 | 4.97E+00 | 2.57E-02 | 1.22E-01 | 1.22E-01 | 2.03E+06 |
| dme-mir-2b-1 | -4.84E-01 | 1.10E+01 | 4.88E+00 | 2.72E-02 | 1.26E-01 | 6.57E-02 | 1.09E+05 |
| dme-mir-1014 | 1.53E+00 | 9.49E-01 | 4.85E+00 | 2.77E-02 | 1.26E-01 | 5.28E-01 | 8.00E+01 |
| dme-mir-303 | 2.26E+00 | -7.32E-01 | 4.75E+00 | 2.92E-02 | 1.30E-01 | 2.90E-01 | 8.00E+00 |
| dme-mir-996 | 4.22E-01 | 1.23E+01 | 4.69E+00 | 3.04E-02 | 1.32E-01 | 5.40E-02 | 2.76E+05 |
| dme-mir-87 | -6.01E-01 | 8.41E+00 | 4.63E+00 | 3.13E-02 | 1.34E-01 | 1.05E-01 | 1.81E+04 |
| dme-mir-997 | 1.82E+00 | -4.22E-01 | 4.12E+00 | 4.25E-02 | 1.78E-01 | 4.31E-01 | 1.60E+01 |
| dme-mir-2b-2 | -4.09E-01 | 1.13E+01 | 4.07E+00 | 4.38E-02 | 1.80E-01 | 5.66E-02 | 1.34E+05 |
| dme-mir-4910 | -9.95E-01 | 1.78E+00 | 3.89E+00 | 4.86E-02 | 1.96E-01 | 2.63E-01 | 1.60E+02 |
| dme-mir-1 | -7.10E-01 | 1.68E+01 | 3.85E+00 | 4.99E-02 | 1.98E-01 | 1.77E-01 | 5.84E+06 |
| dme-mir-190 | -4.45E-01 | 1.06E+01 | 3.75E+00 | 5.28E-02 | 2.06E-01 | 7.23E-02 | 8.24E+04 |
| dme-mir-6-3 | 1.39E+00 | -1.52E-01 | 3.70E+00 | 5.45E-02 | 2.08E-01 | 2.70E-01 | 2.40E+01 |
| dme-mir-283 | 5.51E-01 | 9.47E+00 | 3.61E+00 | 5.75E-02 | 2.16E-01 | 1.19E-01 | 3.88E+04 |
| dme-mir-210 | 5.02E-01 | 1.35E+01 | 3.44E+00 | 6.36E-02 | 2.32E-01 | 1.04E-01 | 6.40E+05 |
| dme-mir-957 | 5.30E-01 | 1.23E+01 | 3.43E+00 | 6.39E-02 | 2.32E-01 | 1.17E-01 | 2.80E+05 |
| dme-mir-975 | 1.39E+00 | -4.50E-01 | 3.24E+00 | 7.17E-02 | 2.56E-01 | 1.46E-01 | 1.50E+01 |
| dme-mir-315 | 5.52E-01 | 1.07E+01 | 3.21E+00 | 7.30E-02 | 2.57E-01 | 1.35E-01 | 9.92E+04 |
| dme-mir-281-2 | 3.81E-01 | 1.02E+01 | 3.14E+00 | 7.62E-02 | 2.64E-01 | 6.53E-02 | 6.46E+04 |
| dme-mir-5 | 9.99E-01 | 5.34E+00 | 3.08E+00 | 7.95E-02 | 2.71E-01 | 4.57E-01 | 2.12E+03 |
| dme-mir-2280 | -1.59E+00 | -5.05E-01 | 3.02E+00 | 8.23E-02 | 2.76E-01 | 3.65E-01 | 1.30E+01 |
| dme-mir-927 | -3.54E-01 | 1.10E+01 | 2.90E+00 | 8.85E-02 | 2.92E-01 | 5.96E-02 | 1.14E+05 |
| dme-mir-2a-1 | -4.10E-01 | 9.55E+00 | 2.80E+00 | 9.44E-02 | 3.07E-01 | 8.22E-02 | 4.01E+04 |
| dme-mir-263b | -3.74E-01 | 1.21E+01 | 2.75E+00 | 9.72E-02 | 3.12E-01 | 7.02E-02 | 2.47E+05 |
| dme-mir-981 | 3.73E-01 | 9.80E+00 | 2.68E+00 | 1.02E-01 | 3.17E-01 | 7.35E-02 | 4.79E+04 |
| dme-mir-988 | -3.99E-01 | 1.01E+01 | 2.68E+00 | 1.02E-01 | 3.17E-01 | 8.14E-02 | 5.97E+04 |
| dme-mir-1000 | 3.85E-01 | 1.15E+01 | 2.55E+00 | 1.10E-01 | 3.38E-01 | 8.23E-02 | 1.54E+05 |
| dme-mir-3645 | -1.69E+00 | -8.26E-01 | 2.52E+00 | 1.12E-01 | 3.40E-01 | 1.84E-01 | 6.00E+00 |
| dme-mir-4945 | -2.16E+00 | 2.58E-01 | 2.48E+00 | 1.16E-01 | 3.45E-01 | 2.04E+00 | 4.20E+01 |
| dme-mir-305 | 3.81E-01 | 1.20E+01 | 2.42E+00 | 1.20E-01 | 3.53E-01 | 8.52E-02 | 2.18E+05 |
| dme-mir-276a | -2.76E-01 | 1.69E+01 | 2.38E+00 | 1.23E-01 | 3.58E-01 | 4.43E-02 | 6.58E+06 |
| dme-mir-2501 | 1.40E+00 | -7.44E-01 | 2.11E+00 | 1.47E-01 | 4.20E-01 | 2.22E-01 | 8.00E+00 |
| dme-mir-281-1 | 3.67E-01 | 8.21E+00 | 2.05E+00 | 1.52E-01 | 4.20E-01 | 9.24E-02 | 1.66E+04 |
| dme-mir-124 | 3.30E-01 | 9.39E+00 | 2.05E+00 | 1.52E-01 | 4.20E-01 | 7.49E-02 | 3.57E+04 |
| dme-mir-318 | -5.56E-01 | 2.50E+00 | 2.02E+00 | 1.56E-01 | 4.20E-01 | 1.65E-01 | 2.74E+02 |
| dme-mir-1011 | 7.25E-01 | 3.72E+00 | 2.01E+00 | 1.56E-01 | 4.20E-01 | 3.55E-01 | 6.99E+02 |
| dme-mir-954 | 3.68E-01 | 6.28E+00 | 2.01E+00 | 1.56E-01 | 4.20E-01 | 9.23E-02 | 4.24E+03 |
| dme-mir-3644 | 1.57E+00 | -8.77E-01 | 1.97E+00 | 1.60E-01 | 4.26E-01 | 3.45E-01 | 5.00E+00 |
| dme-mir-4982 | 1.53E+00 | -8.82E-01 | 1.92E+00 | 1.66E-01 | 4.37E-01 | 3.06E-01 | 5.00E+00 |
| dme-mir-995 | 3.31E-01 | 1.02E+01 | 1.87E+00 | 1.71E-01 | 4.44E-01 | 8.28E-02 | 6.38E+04 |
| dme-mir-1010 | 3.10E-01 | 1.05E+01 | 1.86E+00 | 1.73E-01 | 4.44E-01 | 7.30E-02 | 7.63E+04 |
| dme-mir-307b | -7.45E-01 | 7.91E-01 | 1.80E+00 | 1.80E-01 | 4.56E-01 | 2.49E-01 | 6.70E+01 |
| dme-mir-980 | 4.13E-01 | 3.31E+00 | 1.65E+00 | 1.99E-01 | 4.99E-01 | 1.20E-01 | 4.99E+02 |
| dme-mir-4973 | 6.16E-01 | 1.04E+00 | 1.60E+00 | 2.06E-01 | 5.09E-01 | 1.96E-01 | 8.70E+01 |
| dme-mir-313 | 7.52E-01 | 7.66E-01 | 1.55E+00 | 2.13E-01 | 5.21E-01 | 3.46E-01 | 6.80E+01 |
| dme-mir-317 | -2.29E-01 | 1.78E+01 | 1.43E+00 | 2.32E-01 | 5.60E-01 | 5.11E-02 | 1.16E+07 |
| dme-mir-282 | 3.21E-01 | 8.97E+00 | 1.42E+00 | 2.34E-01 | 5.60E-01 | 1.02E-01 | 2.68E+04 |
| dme-mir-4957 | 7.00E-01 | -6.33E-03 | 1.39E+00 | 2.39E-01 | 5.66E-01 | 1.25E-01 | 2.90E+01 |
| dme-mir-992 | 1.16E+00 | -7.86E-01 | 1.36E+00 | 2.43E-01 | 5.68E-01 | 2.40E-01 | 7.00E+00 |
| dme-mir-2c | 3.58E-01 | 6.14E+00 | 1.35E+00 | 2.45E-01 | 5.68E-01 | 1.31E-01 | 3.93E+03 |
| dme-mir-1005 | 3.78E-01 | 4.93E+00 | 1.29E+00 | 2.56E-01 | 5.81E-01 | 1.49E-01 | 1.65E+03 |
| dme-mir-4 | 7.52E-01 | 3.15E+00 | 1.26E+00 | 2.61E-01 | 5.81E-01 | 6.11E-01 | 4.42E+02 |
| dme-mir-3 | -8.78E-01 | 3.95E+00 | 1.25E+00 | 2.63E-01 | 5.81E-01 | 8.09E-01 | 8.57E+02 |
| dme-mir-4951 | 2.90E-01 | 5.55E+00 | 1.21E+00 | 2.71E-01 | 5.81E-01 | 9.32E-02 | 2.58E+03 |
| dme-mir-11 | -2.50E-01 | 1.23E+01 | 1.21E+00 | 2.71E-01 | 5.81E-01 | 7.14E-02 | 2.72E+05 |
| dme-mir-4943 | 5.53E-01 | 8.37E-01 | 1.20E+00 | 2.73E-01 | 5.81E-01 | 1.98E-01 | 7.30E+01 |
| dme-mir-286 | -4.85E-01 | 3.78E+00 | 1.20E+00 | 2.74E-01 | 5.81E-01 | 2.52E-01 | 7.19E+02 |
| dme-mir-970 | 2.53E-01 | 1.16E+01 | 1.19E+00 | 2.75E-01 | 5.81E-01 | 7.57E-02 | 1.68E+05 |
| dme-mir-955 | 6.09E-01 | 9.02E-01 | 1.19E+00 | 2.75E-01 | 5.81E-01 | 2.82E-01 | 7.60E+01 |
| dme-mir-2497 | 7.93E-01 | 9.17E-01 | 1.17E+00 | 2.80E-01 | 5.81E-01 | 6.24E-01 | 7.80E+01 |
| dme-mir-987 | 2.41E-01 | 1.35E+01 | 1.15E+00 | 2.83E-01 | 5.81E-01 | 7.16E-02 | 6.61E+05 |
| dme-mir-2a-2 | -2.13E-01 | 1.04E+01 | 1.14E+00 | 2.85E-01 | 5.81E-01 | 5.51E-02 | 7.35E+04 |
| dme-mir-969 | -3.14E-01 | 4.95E+00 | 1.13E+00 | 2.87E-01 | 5.81E-01 | 1.13E-01 | 1.68E+03 |
| dme-mir-971 | 2.97E-01 | 5.48E+00 | 1.13E+00 | 2.88E-01 | 5.81E-01 | 1.05E-01 | 2.40E+03 |
| dme-mir-1003 | -3.32E-01 | 6.49E+00 | 1.11E+00 | 2.93E-01 | 5.85E-01 | 1.34E-01 | 4.63E+03 |
| dme-mir-998 | -2.40E-01 | 8.81E+00 | 1.08E+00 | 2.99E-01 | 5.92E-01 | 7.35E-02 | 2.41E+04 |
| dme-mir-4974 | 8.99E-01 | -6.88E-01 | 1.04E+00 | 3.08E-01 | 6.06E-01 | 1.37E-01 | 9.00E+00 |
| dme-mir-4913 | 4.98E-01 | 8.04E-01 | 9.99E-01 | 3.17E-01 | 6.18E-01 | 1.81E-01 | 6.90E+01 |
| dme-mir-193 | 2.17E-01 | 1.10E+01 | 9.52E-01 | 3.29E-01 | 6.35E-01 | 6.96E-02 | 1.10E+05 |
| dme-mir-4919 | 3.75E-01 | 2.53E+00 | 9.32E-01 | 3.34E-01 | 6.40E-01 | 1.69E-01 | 2.80E+02 |
| dme-mir-986 | -2.53E-01 | 1.18E+01 | 8.95E-01 | 3.44E-01 | 6.52E-01 | 9.87E-02 | 1.86E+05 |
| dme-mir-278 | -1.72E-01 | 1.26E+01 | 8.73E-01 | 3.50E-01 | 6.58E-01 | 4.73E-02 | 3.49E+05 |
| dme-mir-1008 | 3.08E-01 | 4.27E+00 | 8.50E-01 | 3.57E-01 | 6.64E-01 | 1.45E-01 | 9.93E+02 |
| dme-mir-4969 | -3.44E-01 | 3.58E+00 | 8.00E-01 | 3.71E-01 | 6.78E-01 | 1.84E-01 | 6.27E+02 |
| dme-mir-2498 | 8.96E-01 | -8.32E-01 | 7.87E-01 | 3.75E-01 | 6.78E-01 | 1.82E-01 | 6.00E+00 |
| dme-mir-4981 | 4.81E-01 | 7.08E-01 | 7.83E-01 | 3.76E-01 | 6.78E-01 | 2.33E-01 | 6.30E+01 |
| dme-mir-307a | 1.98E-01 | 1.10E+01 | 7.82E-01 | 3.76E-01 | 6.78E-01 | 7.03E-02 | 1.05E+05 |
| dme-mir-1004 | -2.32E-01 | 7.57E+00 | 7.53E-01 | 3.85E-01 | 6.80E-01 | 9.78E-02 | 1.03E+04 |
| dme-bantam | -2.16E-01 | 1.45E+01 | 7.45E-01 | 3.88E-01 | 6.80E-01 | 8.70E-02 | 1.26E+06 |
| dme-mir-967 | 3.74E-01 | 1.55E+00 | 7.42E-01 | 3.89E-01 | 6.80E-01 | 1.75E-01 | 1.34E+02 |
| dme-mir-284 | -1.81E-01 | 1.18E+01 | 7.35E-01 | 3.91E-01 | 6.80E-01 | 6.19E-02 | 1.96E+05 |
| dme-mir-4961 | -4.55E-01 | 5.59E-01 | 7.29E-01 | 3.93E-01 | 6.80E-01 | 1.80E-01 | 5.30E+01 |
| dme-mir-2495 | 8.55E-01 | -8.32E-01 | 7.09E-01 | 4.00E-01 | 6.86E-01 | 2.39E-01 | 6.00E+00 |
| dme-mir-275 | 2.27E-01 | 1.05E+01 | 6.85E-01 | 4.08E-01 | 6.95E-01 | 1.06E-01 | 8.10E+04 |
| dme-mir-1013 | 2.61E-01 | 5.71E+00 | 6.56E-01 | 4.18E-01 | 7.06E-01 | 1.42E-01 | 2.94E+03 |
| dme-mir-277 | -1.40E-01 | 1.42E+01 | 6.38E-01 | 4.25E-01 | 7.12E-01 | 4.31E-02 | 9.83E+05 |
| dme-mir-2489 | 3.27E-01 | 1.88E+00 | 6.19E-01 | 4.31E-01 | 7.18E-01 | 1.73E-01 | 1.75E+02 |
| dme-mir-4956 | -5.27E-01 | -3.45E-01 | 5.64E-01 | 4.53E-01 | 7.48E-01 | 1.31E-01 | 1.80E+01 |
| dme-mir-4940 | -5.57E-01 | 2.02E+00 | 5.45E-01 | 4.60E-01 | 7.51E-01 | 7.18E-01 | 1.96E+02 |
| dme-mir-999 | -1.31E-01 | 1.46E+01 | 5.42E-01 | 4.62E-01 | 7.51E-01 | 4.38E-02 | 1.36E+06 |
| dme-mir-13b-2 | 1.91E-01 | 1.22E+01 | 5.28E-01 | 4.68E-01 | 7.54E-01 | 9.71E-02 | 2.64E+05 |
| dme-mir-3641 | -6.14E-01 | -5.35E-01 | 5.11E-01 | 4.75E-01 | 7.54E-01 | 3.06E-01 | 1.30E+01 |
| dme-mir-276b | -1.20E-01 | 1.37E+01 | 5.03E-01 | 4.78E-01 | 7.54E-01 | 4.01E-02 | 7.07E+05 |
| dme-mir-13b-1 | 1.87E-01 | 1.22E+01 | 5.01E-01 | 4.79E-01 | 7.54E-01 | 9.83E-02 | 2.64E+05 |
| dme-mir-4975 | 4.03E-01 | 2.37E-01 | 4.94E-01 | 4.82E-01 | 7.54E-01 | 1.81E-01 | 3.90E+01 |
| dme-mir-8 | 1.67E-01 | 1.51E+01 | 4.88E-01 | 4.85E-01 | 7.54E-01 | 8.06E-02 | 1.87E+06 |
| dme-mir-4909 | 4.47E-01 | -1.06E-01 | 4.72E-01 | 4.92E-01 | 7.54E-01 | 1.78E-01 | 2.50E+01 |
| dme-mir-932 | -1.32E-01 | 1.06E+01 | 4.60E-01 | 4.98E-01 | 7.54E-01 | 5.26E-02 | 8.40E+04 |
| dme-mir-1002 | 5.00E-01 | -1.50E-01 | 4.59E-01 | 4.98E-01 | 7.54E-01 | 3.48E-01 | 2.40E+01 |
| dme-mir-1009 | -2.10E-01 | 4.83E+00 | 4.57E-01 | 4.99E-01 | 7.54E-01 | 1.26E-01 | 1.57E+03 |
| dme-mir-219 | 2.48E-01 | 5.24E+00 | 4.51E-01 | 5.02E-01 | 7.54E-01 | 1.86E-01 | 2.01E+03 |
| dme-mir-2535b | -1.85E-01 | 4.26E+00 | 4.31E-01 | 5.12E-01 | 7.64E-01 | 9.73E-02 | 9.91E+02 |
| dme-mir-263a | 1.47E-01 | 1.44E+01 | 4.15E-01 | 5.19E-01 | 7.66E-01 | 7.33E-02 | 1.17E+06 |
| dme-mir-9a | -1.28E-01 | 1.42E+01 | 4.07E-01 | 5.23E-01 | 7.66E-01 | 5.64E-02 | 9.34E+05 |
| dme-mir-4912 | 7.58E-01 | -9.80E-01 | 4.07E-01 | 5.24E-01 | 7.66E-01 | 3.22E-01 | 3.00E+00 |
| dme-mir-306 | 1.27E-01 | 1.18E+01 | 3.42E-01 | 5.58E-01 | 8.07E-01 | 6.67E-02 | 2.02E+05 |
| dme-mir-308 | 2.41E-01 | 6.60E+00 | 3.39E-01 | 5.60E-01 | 8.07E-01 | 2.40E-01 | 5.15E+03 |
| dme-mir-4984 | 3.31E-01 | 5.81E-01 | 3.31E-01 | 5.65E-01 | 8.07E-01 | 2.60E-01 | 5.50E+01 |
| dme-mir-4952 | 1.77E-01 | 3.87E+00 | 3.25E-01 | 5.68E-01 | 8.07E-01 | 1.18E-01 | 7.62E+02 |
| dme-mir-994 | -2.88E-01 | 1.25E+00 | 3.17E-01 | 5.74E-01 | 8.07E-01 | 2.47E-01 | 1.01E+02 |
| dme-mir-1012 | 1.31E-01 | 9.90E+00 | 3.16E-01 | 5.74E-01 | 8.07E-01 | 7.64E-02 | 4.98E+04 |
| dme-mir-2493 | -3.23E-01 | 1.31E-01 | 3.05E-01 | 5.81E-01 | 8.09E-01 | 1.56E-01 | 3.40E+01 |
| dme-mir-1016 | -1.99E-01 | 2.49E+00 | 3.02E-01 | 5.83E-01 | 8.09E-01 | 1.37E-01 | 2.70E+02 |
| dme-mir-4986 | -4.30E-01 | -4.50E-01 | 2.95E-01 | 5.87E-01 | 8.10E-01 | 2.53E-01 | 1.50E+01 |
| dme-let-7 | 8.95E-02 | 1.53E+01 | 2.66E-01 | 6.06E-01 | 8.26E-01 | 4.23E-02 | 2.08E+06 |
| dme-mir-4946 | -6.07E-01 | -9.76E-01 | 2.65E-01 | 6.06E-01 | 8.26E-01 | 3.22E-01 | 3.00E+00 |
| dme-mir-4918 | -4.22E-01 | -6.46E-01 | 2.47E-01 | 6.19E-01 | 8.33E-01 | 1.36E-01 | 1.00E+01 |
| dme-mir-133 | -9.74E-02 | 1.21E+01 | 2.47E-01 | 6.19E-01 | 8.33E-01 | 5.35E-02 | 2.35E+05 |
| dme-mir-2496 | -4.32E-01 | -2.42E-01 | 2.42E-01 | 6.23E-01 | 8.33E-01 | 6.31E-01 | 2.10E+01 |
| dme-mir-4944 | -5.50E-01 | -9.84E-01 | 2.32E-01 | 6.30E-01 | 8.38E-01 | 3.22E-01 | 3.00E+00 |
| dme-mir-7 | -1.29E-01 | 1.53E+01 | 2.20E-01 | 6.39E-01 | 8.44E-01 | 1.05E-01 | 2.34E+06 |
| dme-mir-6-2 | 4.18E-01 | 3.03E-02 | 1.97E-01 | 6.57E-01 | 8.55E-01 | 9.37E-01 | 3.00E+01 |
| dme-mir-13a | -1.34E-01 | 8.06E+00 | 1.96E-01 | 6.58E-01 | 8.55E-01 | 1.28E-01 | 1.41E+04 |
| dme-mir-274 | 9.96E-02 | 1.47E+01 | 1.93E-01 | 6.60E-01 | 8.55E-01 | 7.22E-02 | 1.40E+06 |
| dme-mir-316 | 1.13E-01 | 6.40E+00 | 1.89E-01 | 6.63E-01 | 8.55E-01 | 9.21E-02 | 4.54E+03 |
| dme-mir-252 | -7.78E-02 | 1.38E+01 | 1.85E-01 | 6.67E-01 | 8.55E-01 | 4.57E-02 | 8.06E+05 |
| dme-mir-6-1 | -3.16E-01 | -3.52E-01 | 1.74E-01 | 6.77E-01 | 8.59E-01 | 3.17E-01 | 1.80E+01 |
| dme-mir-2283 | -2.30E-01 | 1.89E-01 | 1.70E-01 | 6.80E-01 | 8.59E-01 | 1.45E-01 | 3.70E+01 |
| dme-mir-4987 | -4.21E-01 | -7.37E-01 | 1.68E-01 | 6.82E-01 | 8.59E-01 | 6.70E-01 | 8.00E+00 |
| dme-mir-4947 | -2.93E-01 | -4.07E-01 | 1.46E-01 | 7.02E-01 | 8.77E-01 | 2.10E-01 | 1.60E+01 |
| dme-mir-79 | -8.78E-02 | 7.90E+00 | 1.42E-01 | 7.06E-01 | 8.77E-01 | 7.46E-02 | 1.27E+04 |
| dme-mir-4968 | -1.46E-01 | 2.07E+00 | 1.41E-01 | 7.08E-01 | 8.77E-01 | 1.50E-01 | 2.02E+02 |
| dme-mir-990 | -1.06E-01 | 5.01E+00 | 1.25E-01 | 7.23E-01 | 8.91E-01 | 1.18E-01 | 1.75E+03 |
| dme-mir-4949 | -2.18E-01 | -9.14E-02 | 1.19E-01 | 7.30E-01 | 8.94E-01 | 1.40E-01 | 2.60E+01 |
| dme-mir-4953 | -3.83E-01 | -8.72E-01 | 1.13E-01 | 7.37E-01 | 8.98E-01 | 6.60E-01 | 5.00E+00 |
| dme-mir-2492 | -3.48E-01 | -8.77E-01 | 1.07E-01 | 7.44E-01 | 9.01E-01 | 3.76E-01 | 5.00E+00 |
| dme-mir-4971 | -2.44E-01 | -4.11E-01 | 9.37E-02 | 7.60E-01 | 9.08E-01 | 3.01E-01 | 1.60E+01 |
| dme-mir-2282 | -2.32E-01 | -3.09E-01 | 9.24E-02 | 7.61E-01 | 9.08E-01 | 3.36E-01 | 1.90E+01 |
| dme-mir-309 | 3.03E-01 | 2.29E-01 | 8.82E-02 | 7.66E-01 | 9.08E-01 | 1.23E+00 | 3.90E+01 |
| dme-mir-4980 | -2.95E-01 | -7.85E-01 | 8.77E-02 | 7.67E-01 | 9.08E-01 | 2.83E-01 | 7.00E+00 |
| dme-mir-279 | 7.32E-02 | 1.05E+01 | 8.50E-02 | 7.71E-01 | 9.08E-01 | 8.84E-02 | 7.55E+04 |
| dme-mir-4958 | 1.36E-01 | 1.48E+00 | 8.15E-02 | 7.75E-01 | 9.09E-01 | 2.20E-01 | 1.25E+02 |

